# A Novel Computational Approach for the Mining of Signature Pathways Using Species Co-occurrence Networks in Gut Microbiomes

**DOI:** 10.1101/2023.08.28.555189

**Authors:** Suyeon Kim, Ishwor Thapa, Hesham Ali

## Abstract

Advances in metagenome sequencing data continue to enable new methods for analyzing biological systems. When handling microbial profile data, metagenome sequencing has proven to be far more comprehensive than traditional methods such as 16s rRNA data, which rely on partial sequences. Microbial community profiling can be used to obtain key biological signals that pave the way for better and accurate understanding of complex systems that are critical for advancing biomedical research and healthcare. There have been few attempts to uncover microbial community associations with certain health conditions. However, such attempts have mostly used partial or incomplete data to accurately capture those associations. This study introduces a novel computational approach for the identification of co-occurring microbial communities using the abundance and functional roles of species-level microbiome data. The proposed approach is then used to identify signature pathways associated with inflammatory bowel disease (IBD). Furthermore, we developed a computational pipeline to identify microbial species co-occurrences from metagenome data. When comparing IBD to a control group, we show that co-occurring communities of species are enriched for potential pathways. We also show that the identified co-occurring microbial species operate as a community to facilitate pathway enrichment. The obtained findings suggest that the proposed network model, along with the computational pipeline, provide a valuable analytical tool to analyze complex biological systems and extract pathway signatures that classify or diagnose certain health conditions.

**CCS CONCEPTS:** **• Applied computing** → **Biological networks**; **Bioinformatics**; **Systems biology**.

**ACM Reference Format:** Suyeon Kim, Ishwor Thapa, and Hesham Ali. 2023. A Novel Computational Approach for the Mining of Signature Pathways Using Species Co-occurrence Networks in Gut Microbiomes. In *Proceedings of ACM Conference (Conference’17)*. ACM, New York, NY, USA, 10 pages. https://doi.org/10.1145/nnnnnnn.nnnnnnn

## 1 INTRODUCTION

Recent advances in high throughput data have revolutionized the way we understand the role of microbiomes in various environments, including the microbiome communities in the human body. The potential for microbiome-based interventions increases as we improve our understanding of microbiome associated health conditions [12]. The characterization of microbial roles in host health and disease is therefore essential for achieving non-invasive, personalized, and accelerated treatment options. Microbial communities vary depending on their age, geography, and population. Given the complex variation in host ecosystems, there is no gold standard profile to identify microbial community patterns and their underlying roles in human health and disease [4].

Despite recent progress in microbiome research, we have yet to identify specific microbial patterns associated with host health phenotypes. This is mainly due to microbial heterogeneity and diverse functional roles. In addition, various computational tools exist for the identification of differentially abundant microbes from abundance data and for microbiome network-based analysis, including NetCoMi [27] and iNAP [8]. However, their are very few computational pipelines for the identification of signature pathways based on co-occurring microbial species.

Different groups in microbial communities may have extremely similar biological functions, especially evidenced in healthy individuals [4, 6]. These findings suggest the possibility of specific microbiome profiles that are associated with healthy phenotypes. However, it is clearly a challenging research question to identify such profiles. An even more challenging question would be to identify microbial characteristics or profiles that are associated with certain unhealthy or disease phenotypes. Establishing such associations would be extremely helpful in early diagnosis of some illnesses, and even more important in personalized treatments. Other issues need to be considered when answering these research questions.

We have the basic notion about levels of certain phyla or ratios of different phyla to be higher or lower for a given phenotype, but these expectations are not always consistent across different studies with different age groups and various ethnicities. For example, when looking at obese patients, the ratio of Firmicutes to Bacteriodes is higher in Caucasian versus Asian populations [21].

Additionally, bioinformatics tools like the HMP Unified Metabolic Analysis Network (HUMAnN) are widely used to generate taxonomic, functional, and strain-level profiles from raw metagenomic sequencing data [1]. While the output of such tools can be directly interpreted with basic downstream analyses, they may not be ideal for distinguishing microbial community profiles (composition and inter-relationships) in disease states versus healthy states. To this end, shotgun metagenomic sequencing data are now publicly available and can be readily obtained from multiple independent studies around the world. Pooling such datasets, while challenging, would provide a reliable source to identify robust and reproducible microbial signatures that are consistent across many different studies.

In this study, we propose a novel computational pipeline to identify microbiome-based functional enrichment patterns associated with host disease states. The proposed pipeline focuses on mining a high-level summary of the enriched pathways underlying given microbial communities that are associated with host health and disease states. This pipeline is designed to allow researchers to analyze microbial communities at different granularity levels: a single microbe, microbial sub-communities, and overall microbial network structures. When comparing microbial communities in healthy versus disease states, it is important to answer several research questions: What bacterial species are central in these communities? How do community networks differ across phenotypes? How conserved are these communities across related datasets? What sub-group of bacterial species (or clusters) in these communities are functionally enriched in a specific disease state?

Our pipeline offers flexible functionality for performing differential abundance analysis on any meta-omics datasets. The input data is used to construct and analyze co-occurrence networks in a comprehensive manner. While this study is focused on samples for IBD groups, the overarching goal of the study is to develop a computational pipeline with generalizabiltiy beyond inflammatory bowel disease (IBD). In this study, we used microbiome samples for two inflammatory bowel disease (IBD) groups, consisting of Crohn’s disease (CD) and Ulcerative colitis (UC) from two independent studies.

## 2 PROBLEM DESCRIPTION AND PROPOSED APPROACH

Every time biomedical technologies provide access to new types of biological data, researchers tend to use similar approaches to address similar problems. Would it possible for the new available data to provide new biological signals that can be used for classification purposes? Can we use the new data types to profile groups with common phenotypes? Would the biological signals associated with the new data be robust enough to be used for the purpose of disease diagnosis and/or the assessment of different treatments for certain health conditions. New studies are needed to explore such potential. Similar patterns have occurred with the availability of data for gene expression, genetic variants, and protein interactions. Now with the availability of microbiome data, similar questions are emerging.

The availability of microbiome data provides the possibility of obtaining new characterizing signals that can be used for classification purposes. Hence, the goal of this study is to establish helpful associations between the profiles of gut microbiomes and health features. It would have been straightforward to establish such associations by using the abundance levels of species contained in microbiomes, unfortunately, this may not always be possible. Microbiome species play different roles at different situations and similar biological functions can be performed by different species. Therefore, we need to build our approach around the functional roles of the microbiome species as well as their abundance levels.

In this work, the main goal is to develop a computational pipeline to identify possible associations between gut microbiomes and phenotypes related to specific diseases. We explore microbiome profiles based on the abundances and co-occurrences of their species along with their functional roles in known pathways. We test the viability of identifying characterizing pathways enriched by high co-occurring microbiome species for subjects with conditions such as CD and UC.

We also try to address the issue of robustness in this study. Identifying phenotypic-level associations from metagenome sequencing is challenging because it contains DNA sequences from millions of microbes that are collectively working to achieve specific biological functions. Researchers traditionally investigate a single species at a time, which is insufficient for understanding its contribution to a disease phenotype. However, identifying unique microbial signatures associated with host health conditions from multiple datasets is not trivial due to variability in samples across those studies. Our approach captures co-occurring species to efficiently identify robust pathway signatures in different disease conditions, all while comparing multiple datasets concurrently.

## 3 METHODS

Due to many confounding factors, the role of a microbial community associated with a specific host phenotype is not reproducible across multiple datasets. To overcome this challenge and identify microbial species and communities associated with pathways conserved across different datasets, we introduce a novel computational pipeline that contains 4 major steps: (1) construct a species co-occurrence network from filtered samples; (2) Analyze the species co-occurrence network at three granularity levels (single microbe, microbial sub-community, and overall microbial network structure); (3) Perform pathway enrichment analysis; (4) Compare enriched pathways between two IBD datasets in each sample from healthy/disease states.

To achieve this, we combine enriched pathways across the afore-mentioned three analyses to identify robust enriched pathways with contributing species, thereby indicating possible functional markers associated with IBDs. An overview of the computational pipeline is shown in Figure 1.

**Figure 1:**
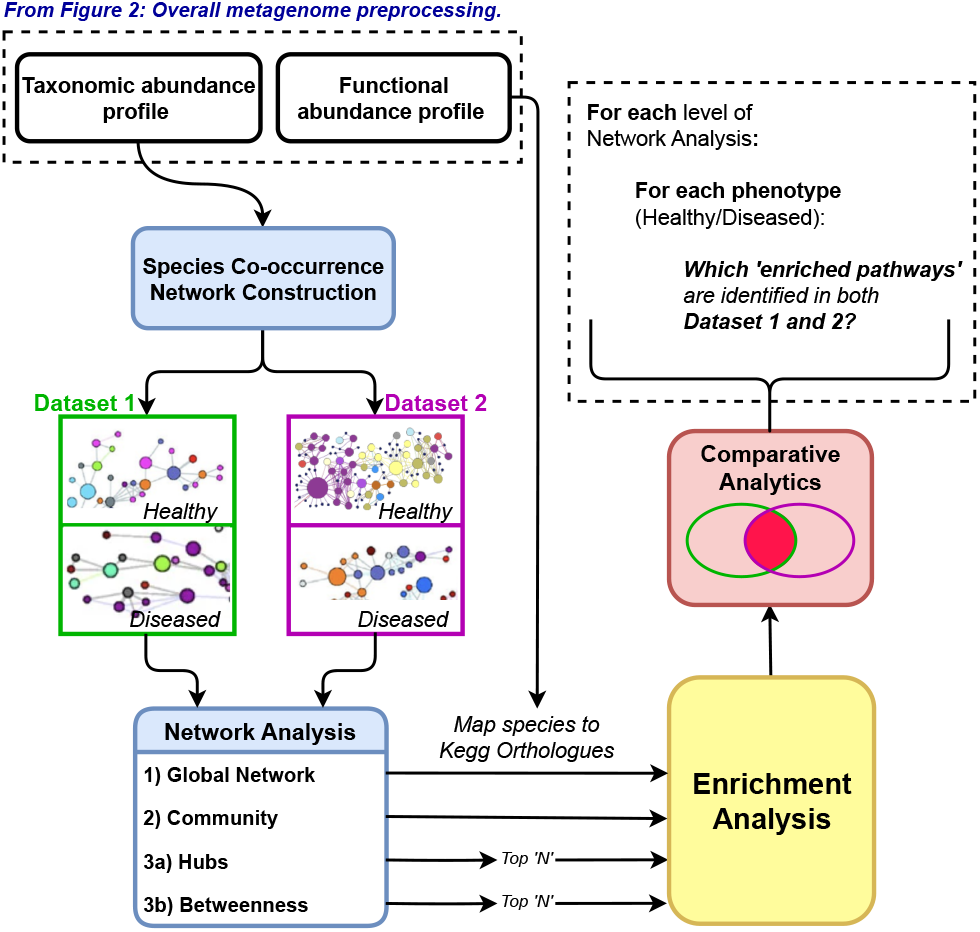
Novel computational pipeline for mining signature pathways that are associated with host health conditions:(A) Construction of species co-occurrence network using taxonomic profiles; (B) Microbial species co-occurrence network analysis; (C) Enrichment analysis; (D) Identification of robust enriched pathway.

### 3.1 Data Pre-processing

As a case study to show the utility of this computational pipeline, we obtained two publicly available human stool metagenome datasets and their metadata. Raw metagenomic sequences of two published inflammatory bowel disease (IBD) studies were obtained from NCBI Sequenced Read Archive PRJNA398089 [18] and PRJNA400072 [10].

#### Sample selection criteria

Samples were removed based on several exclusion criteria. We excluded samples pertaining to medication interventions (i.e., antibiotics, immunosuppressant) and participants who were younger than 18 or older than 65 years of age. Furthermore, duplicate samples IDs were removed.

The study from Franzosa et al. [10] employed cross-sectional data from subjects. Since the study from Lloyd-Price et al. [18] employed longitudinal sampling of subjects, only samples from the subject’s initial doctor visit were included.

##### 3.1.1 Profiling the abundance of microbiome and other molecular functions

Metagenomic samples were taken from two IBD cohorts containing Crohn’s disease (CD) and ulcerative colitis (UC), as well as non-IBD controls. Analysis was performed using HUMAnN [1]. A summary of sample sizes after data pre-processing is shown in Table 1.

**Table 1:** Overview of the number of samples used in the study.

| Dataset 1 |  |  |  |
| --- | --- | --- | --- |
| non-IBD controls | CD | UC | Total |
| 13 | 26 | 18 | 57 |
| Dataset 2 |  |  |  |
| non-IBD controls | CD | UC | Total |
| 54 | 69 | 48 | 171 |

MetaPhlAn version 3.1.0, which can provide pan-microbial (bacterial, archaeal, viral, and eukaryotic) profiling, was used to generate taxonomic profiles of shotgun metagenomes from 57 and 171 samples in dataset 1 and dataset 2, respectively. Functional profiling was performed by HUMAnN version 3.1.1. The output of this tool includes gene-families abundance profiles (UniRef90s), which can be summarized as KEGG Orthologues (KOs).

##### 3.1.2 Species and samples filtering based on taxonomic profiles

After taxonomic profiling, we removed archaea, eukaryota, and zero prevalance species across non-IBD control, CD, and UC phenotypes. The zero prevalence species are those that are not present in any of the samples. Note that the focus of this study is to construct species-level co-occurrence, as the species level taxon provides comprehensive microbiome information. With the species abundance data, we measured Shannon diversity index to calculate species diversity and richness using the R ‘vegan’ package [25]. Shannon diversity index is a widely used method for measuring biological diversity [2]. Samples with zero Shannon diversity index were excluded. A schematic overview of metagenome preprocessing is shown in Figure2.

**Figure 2:**
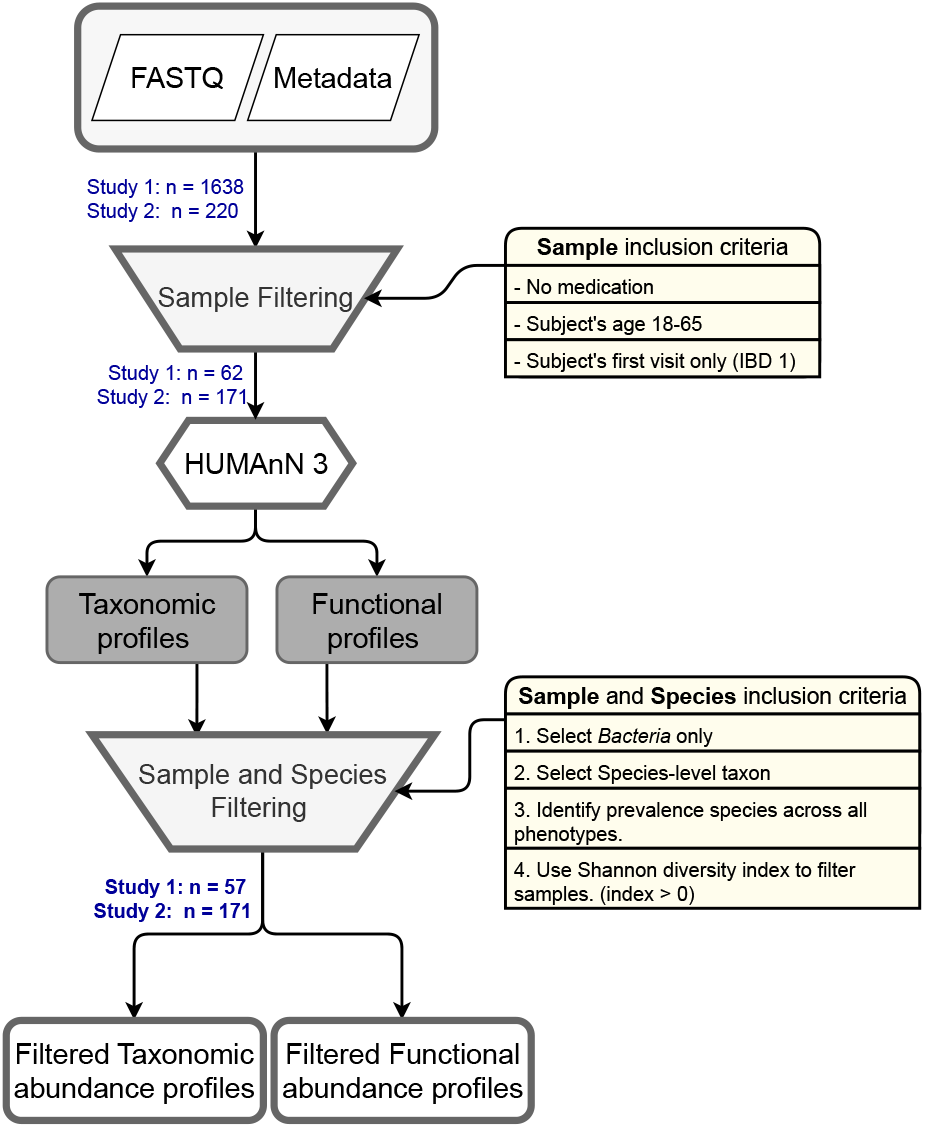
Flowchart depicting a microbial species profile from multiple studies filtering process. Rounded rectangles indicate inputs and outputs. A hexagon indicates a process using a software tool and trapezoids indicate sample and species filtering processes.

### 3.2 Creating the species co-occurrence network

After pre-processsing the species-level abundance and subsequent filtration of species and samples, the species relative abundance matrix was converted into species presence/absence matrix. Abundance values greater than zero were considered “present”. Based on this matrix, co-occurrence probabilities at the species level were measured in each phenotype. To accomplish this, we utilized the probabilistic model of species co-occurrence from the ‘*cooccur*’ R package [11]. The co-occurrence probabilities for each species pair were retrieved as a table containing the pair information, observed number of samples having both species, and the probability of both species occurring in samples from a phenotype. The significance levels of positive and negative co-occurrence patterns were obtained by measuring how greater or less the probabilities of those co-occurring species were than the observed frequencies. Then, from all to all species pairs, we extracted significantly positive co-occurrence pairs with ‘*p*_*gt* ′ *<* 0.05. For the significant positive species pairs, we constructed the unweighted species co-occurrence network, where the species are represented as nodes and edges are drawn if two species are significantly co-occurring in that phenotype within each dataset.

### 3.3 Network Analysis

Next we study different network properties in these co-occurrence networks from all phentoypes in two datasets. Network properties can be evaluated at multiple granularity levels. For example, an element level with a focus on a node or an edge, community level with a focus on a group of nodes and edges, and finally the complete networks. In this study, we examined the species co-occurrence networks at all three levels. These three levels of analyses were performed on species co-occurrence networks from each phenotype in both IBD datasets as described below:

#### 3.3.1 Global network analysis

We analyzed the overall species co-occurrence network from the non-IBD control, CD, and UC groups. For each group, we performed functional enrichment analysis based on the total number of species captured from these co-occurrence networks. Enriched pathways for each diseased group were compared against the non-IBD control. We evaluated the pathways that were consistently enriched across multiple datasets.

#### 3.3.2 Community-level analysis

To identify groups of species that were highly connected between each other, we performed the Leiden community detection algorithm to detect species-communities in the networks from each phenotype. We used the Constant Potts Model (CPM) as the objective function. The value of the resolution parameter is based on the ratio of the quartile value of strength of the nodes and the total number of nodes as shown in the igraph package in R [5]. Functional enrichment of each cluster (community) of bacterial species were compared with the notion that bacterial communities work together to achieve a function. Across two datasets, we identified the functions that were consistently enriched within the same phenotype.

#### 3.3.3 Key element-level analysis

Basic descriptive network properties of the species co-occurrence networks in non-IBD control, CD, and UC were analyzed using the *igraph* R package [5]. We measured centrality of the network to measure topological importance of nodes within the network. Specifically, hub and betweenness centrality were quantified. ‘Hub’ measures the degree of connectedness of a node to all other nodes within a network and can be inferred as how central the node is based on the eigenvector centrality. ‘Betweenness centrality’ of a node is a measurement of how many shortest paths within the network pass through the node, which is critical to maintaining the connectedness of that network. We obtained the top 5 species nodes with hub and betweenness centrality, respectively. In this case, the selection of threshold did not impact on the result. Further, we characterize the roles of these central nodes by performing functional enrichment analysis and compare those across datasets.

### 3.4 Functional enrichment analysis

#### 3.4.1 Mapping Species-to-KO

The obtained list of species from the aforementioned analyses from three levels were then mapped to their KEGG Ortholog identifiers (KO). This mapping step requires KO abundance profiles generated by the HUMAnN pipeline. The abundance profiles from the HUMAnN pipeline contain abundances of KOs stratified by each species and an aggregate abundance value for that KO. We mapped our species of interest to KOs that they contribute to. Next, this list of KOs was used to perform pathway enrichment analysis.

#### 3.4.2 Pathway enrichment analysis

After mapping species with their KO information, we performed KEGG pathway enrichment analysis with the *enrichKO* function in the *MicrobiomeProfiler* package in R [3]. This function performs hypergeometric test to find enrichment for each pathway by utilizing the number of KOs from our species of interest, the KOs annotated for a pathway and all the KOs in the background (universe). The result table contains the pathway information, gene ratio, adjusted p-value, geneIDs for each pathway.

### 3.5 Comparative Analyses

For each dataset and at different network level analyses, we compared the corresponding lists of enriched pathways between disease phenotype and healthy phenotype. Subsequently, the pathways unique to disease conditions are compared across two datasets. These identified unique pathways across datasets indicate possible functional markers associated with the disease phenotype. We visualized those identified common pathways associated with diseased phenotypes to determine which genes from the co-occurring species are contributing to those pathways. We generated this pathway of interest graph by specifying the list of gene IDs (KO-IDs) and pathway ID using the ‘*pathview*’ function [19].

## 4 RESULTS

### 4.1 Global comparison of co-occurrence networks in different health conditions

The total number of significant positive species co-occurrences from non-IBD control, CD, and UC are shown in Table 2. Co-occurrence networks in dataset 2 are larger than in dataset 1 across phenotypes. Between different phenotypes, the co-occurrence network from CD is always bigger than the remaining two.

**Table 2:** Summary of significant co-occurrence of species for non-IBD control, CD, and UC in each dataset.

| Dataset 1 |  |  |  |
| --- | --- | --- | --- |
|  | non-IBD controls | CD | UC |
| p-value < 0.05 | 522 | 721 | 329 |
| Dataset 2 |  |  |  |
|  | non-IBD controls | CD | UC |
| p-value < 0.05 | 2743 | 3722 | 3987 |

#### 4.1.1 Pathways enriched in global species co-occurrence networks for diseased groups across datasets

After pathway enrichment, we compared the enriched pathways in diseased groups (CD and UC) against the non-IBD control and identified common pathways across two IBD datasets (as shown in Table 3). With a significance less than 0.05, the *ABC transporter pathway (map02010)* was enriched in the co-occurring species from CD in both datasets. Similarly, the *Cysteine and methionine metabolism pathway (map00270)* was common to UC in both datasets. The total number of co-occurring species for those enriched pathways in CD and UC are described in Table 3.

**Table 3:** Result of common enriched pathways in CD and UC from global co-occurrence network.

| Phenotype | Enriched pathways | No. co-occurrence species |
| --- | --- | --- |
| CD | ABC transporters (map02010) | 165 |
| UC | Cysteine and methionine metabolism (map00270) | 144 |

### 4.2 Enriched pathways based on community species

The co-occurring species in a community may indicate similar functional capability. We investigated this through clustering and pathway enrichment analysis. For each co-occurrence network, we extracted clusters and identified the membership of the clusters. In CD, we detected 10 clusters in IBD dataset 1 and 55 clusters in IBD dataset 2. UC also contained 7 and 61 clusters in these two datasets, respectively. Then, pathway enrichment of each cluster of species was compared across the datasets. We found 3 enriched pathways for CD and 1 enriched pathway for UC, showing that these pathways were consistently identified between two IBD datasets. We also summarized the number of co-occurring species for each cluster that were found in the enriched pathways associated with diseased groups (as shown in Table 4). We found a sulfur relay system (map04122) was enriched in multiple clusters for both CD and UC. We also found that differences in enriched pathways distinguished CD: *ABC transporter (map02010)* and *Two-component system (map02020)*. Note that a ABC transporter pathway was also enriched in global species co-occurrence networks in CD. Overall, these results showed that these co-occurring species in each cluster are highly enriched for distinct pathways for CD and UC, suggesting these species need further study.

**Table 4:** Table showing the common enriched pathways for each diseased group across two IBD datasets and the number of co-occurrence species for each cluster belonging to the enriched pathway.

| Phenotype | List of enriched pathways | No. Sp in IBD 1 | No. Sp in IBD 2 |
| --- | --- | --- | --- |
| CD | ABC transporter (map02010) | 10 (CL 5) | 129 (CL 2) |
|  | Two-component system (map02020) | 7 (CL 5) | 119 (CL 2) |
|  | Sulfur relay system (map04122) | 4 (CL 5) | 1 (CL 54) |
| UC | Sulfur relay system (map04122) | 3 (CL 4) | 1 (CL 57) |

A visualization of identified clusters enriched in sulfur relay pathways in CD and UC are shown in Figure 3. A total of 15 bacterial species were populated in cluster 5 and enriched for the sulfur relay pathway. We observed 4 species that were present in co-occurring networks in CD from dataset 1. For UC in dataset 1, we found 4 bacterial species were present in cluster 54, and among these 4, KOs from 3 species were enriched in the sulfur replay pathways. We also observed clusters in CD and UC from dataset 2, which are enriched for sulfur relay system. The *Haemophilus sp HMSC71H05* species alone was identified in both cases.

**Figure 3:**
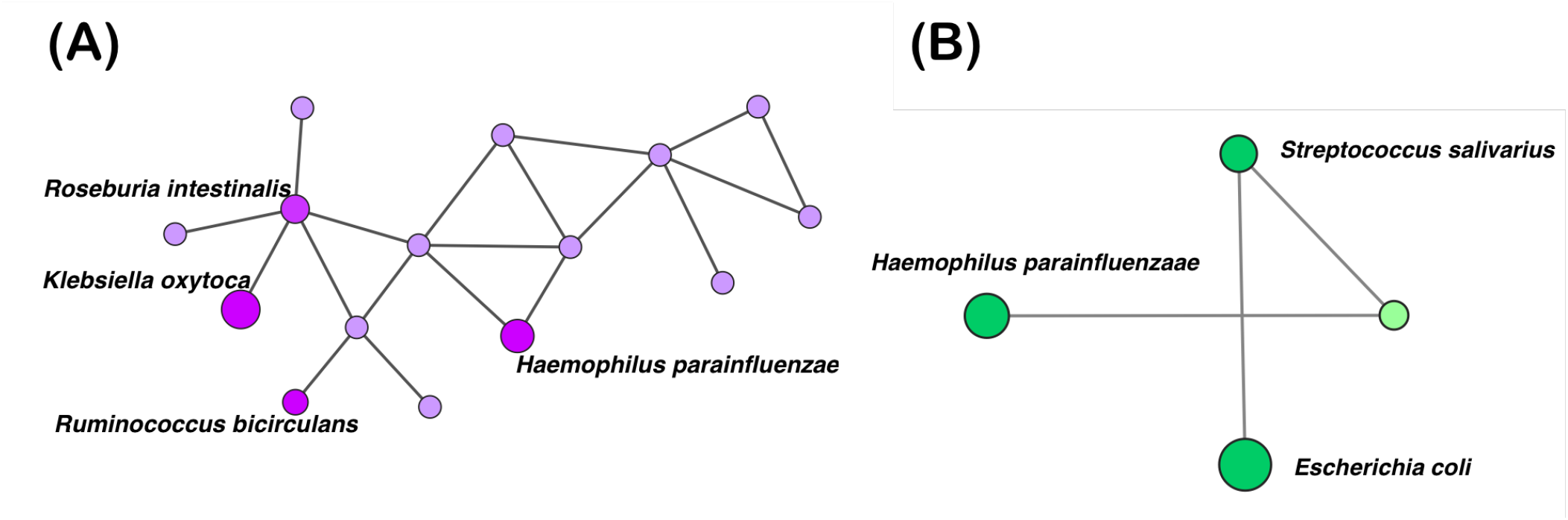
Sulfur relay system pathway for each microbial community in CD dataset 1 (A) and UC dataset 1 group (B). Nodes represent species. An edge is drawn between nodes if two species co-occur together. Nodes are colored by their diseased group (purple indicates CD and green indicates UC). The size of the nodes corresponds to the number of KOs each bacteria contributes in the sulfur relay pathway. Nodes are colored brighter if they were identified in our results from Table 4.

### 4.3 Co-occurring species feature sulfur relay system pathway

Additionally, we visualized the pathway graphs in order to investigate which co-occurring species’s KOs (genes) are contributing to the commonly enriched pathway (Fig 4). For example, the sulfur relay systems (map04122) pathway from CD was further analyzed with the KOs attributable to species (highlighted in red) (see Fig 4). A total of 29 genes are encoding the sulfur relay system pathway while 20 genes were identified in co-occurring species (these genes were highlighted). The co-occurring species counts for sulfur relay system pathway cluster (CL 5) in CD and UC between two datasets, respectively, are shown in Table 4: 4 co-occurring species in CD 1 and 1 species in CD 2. Similarly, 3 co-occurring species in UC 1 and 1 species in UC 2. The *Haemophilus parainfluenzae* and *Haemophilus sp HMSC71H05* species were identified in both CD and UC. The *Klebsiella oxytoca* species has the most number of KOs in this pathway.

**Figure 4:**
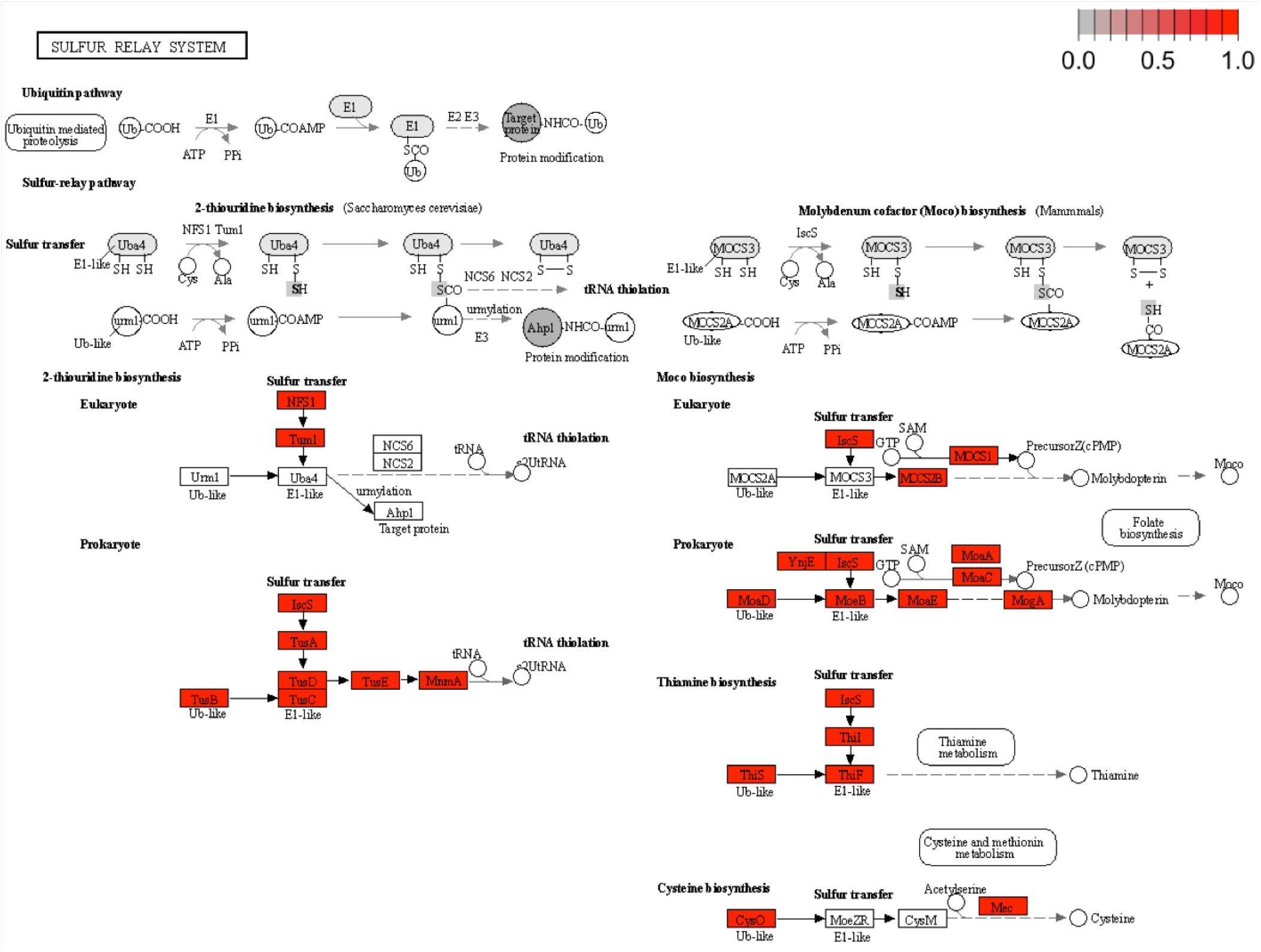
Sulfur relay system pathway with KOs(gene) from co-occurring species in CD. KOs from co-occurring species are highlighted in the color red.

We quantified the presence of KOs in each species (Table 5) and found that our identified community of bacterial species have KOs, contributing to a majority of the prokaryotic sulfur relay system pathway. Note that not a single bacterium is able to encode KOs to complete this pathway by itself.

**Table 5:** List of co-occurring species and count of KOs contributing to the *Sulfur Relay System* pathway.

| Phenotype | Name of species | Number of KOs |
| --- | --- | --- |
| CD 1 | *Haemophilus parainfluenzae* | 9/29 |
|  | *Klebsiella oxytoca* | 18/29 |
|  | *Roseburia intestinalis* | 6/29 |
|  | *Ruminococcus bicirculans* | 1/29 |
| CD 2 | *Haemophilus sp HMSC71H05* | 8/29 |
| UC 1 | *Escherichia coli* | 15/29 |
|  | *Haemophilus parainfluenzae* | 9/29 |
|  | *Streptococcus salivarius* | 1/29 |
| UC 2 | *Haemophilus sp HMSC71H05* | 8/29 |

### 4.4 Enriched pathways based on centrality

We identified KOs from the top 5 species by two centrality measures (hub and betweenness) for enrichment in CD and UC from both IBD datasets. Enriched pathways for CD and UC were compared to identify pathways that are consistently enriched in each phentoype across datasets. We observed that the top 5 hub species, indicating are highly connected within the non-IBD controls, CD, and UC co-occurrence networks from both IBD datasets. We compared enrichment results for these top 5 species and their KOs for CD and UC against non-IBD controls. We also found that list of species with betweenness centrality for the UC in dataset 1 (*Alistipes shahii, Alistipes finegoldii, Oscillibacter sp 57 20, Collinsella aerofaciens*, and *Ruminococcus torques*) are different from dataset 2 (*Asaccharobacter celatus, Clostridium leptum, Eubacterium siraeum, Firmicutes bacterium CAG 103*, and *Ruminococcus torques*). The common enriched pathways for the top 5 hub species from CD and UC in two datasets are shown in Table 4. For CD, we observed the top 5 species *Alistipes shahii, Bacteroides thetaiotaomicron, Coprococcus comes, Odoribacter splanchnicus*, and *Parabacteroides merdae* based on top hub scores in dataset 1 and also in dataset 2, including *Alistipes finegoldii, Asaccharobacter celatus, Blautia obeum, Eubacterium hallii*, and *Lachnospiraceae bacterium OF09 33XD*. The most enriched pathways for CD were *pyrimidine metabolism (map0024)* and *streptomycin biosynthesis (map00521)*, which was not identified in UC. For UC, *lysine biosynthesis (map00300)* and *peptidoglycan biosynthesis (map00550)* pathways were the uniquely enriched pathways. Similarly, we also examined the top 5 species with high betweenness centrality in CD and UC co-occurrence networks. Enrichment results for CD and UC versus non-IBD controls are shown in Table 5. While we found 4 enriched pathways in the UC, no enriched pathways were found in CD.

## 5 DISCUSSION

In this work, we propose a computational pipeline to identify a microbiome-based functional enrichment pattern in host disease states. The main goal of the proposed pipeline is to identify microbial communities that are collectively working to achieve specific biological functions. It has been reported in several studies that co-occurring microbial species with high frequency may be indicative of biological functions governing community structure [17, 20]. Once identified, the pathway enrichment analysis further perform to test the biological significance and identified microbial communities that are collectively enriched in unique pathways associated with health/disease phenotype. The results presented here illustrate the need for augmenting species co-occurrence network with functional-level analysis to identify health/disease associated signatures.

While numerous network inference methods to infer microbial co-occurrences from microbiome abundance datasets are available, it is still difficult to identify true microbial co-occurrences within a complex microbial ecosystem. Therefore, we performed another inference method, SPIEC-EASI [15] to infer the species co-occurrence network, and compared it to the probabilistic approach using the ‘cooccur’ R package. We found that not all highly inferred microbial species interactions with SPIEC-EASI were captured using ‘cooccur’. Upon manual inspection of some of the highly inferred interactions from SPIEC-EASI, we observed that their abundance values were almost zero across all samples in that phenotype and were highly correlated. Due to these spurious correlations in this approach, we believe the probabilistic method from ‘cooccur’ is more suitable in this case.

We analyzed the network properties in these co-occurrence networks at multiple granularity levels. Interestingly, we found that most of the central nodes (based on hub and betweenness measures) appeared in only one of the highly connected clusters obtained from the community level analysis. We note that different pathways from significant species enrichment were identified at different granularity levels. However, all of these pathways were relevant to IBD. This suggests that analysis at multiple granularity levels provided a unique opportunity to identify significant pathways relevant to IBD.

As seen in Table 2, we observed that there were more significant co-occurrences of species in the second dataset. This could be due to the larger number of samples in this study. We first compared the similarity of the species co-occurrence networks for three phenotypes (CD-CD, UC-UC, and control-control) across datasets. Jaccard index was used to determine how many of species in the network for dataset 1 overlapped with that of dataset2. For each phenotype, we expected the Jaccard index score for each comparison to be high; however, this is not the case. The Jaccard index scores for control-control, CD-CD and UC-UC were 0.27, 0.42, and 0.32, respectively. Zelezniak et al. [30] noted that if communities involve phylogenetically related species, then they are likely to be functionally dependent. For example, if ‘Genus A - Species B’ is observed in dataset 1 and ‘Genus A - Species C’ is detected in dataset 2, the Species C in dataset 2 may be functionally similar to Species B in dataset 1.

Given this knowledge, co-occurring species pairs may not be reproducible across independent datasets. However, species level analysis can be useful to understand biological insights in depth. Therefore, we focus on pathway-based signatures between the IBD dataset rather the species signatures. The two enriched pathways in each diseased group at the global network analysis is shown in Table 3. The *ABC transporters (map02010)*, an enriched pathway in CD phenotype, is one of the membrane transporters involving uptake of a variety of small molecules such as nutrients, ions, and drugs [7, 13]. Previous studies have shown associations between transport alterations and IBD pathophysiology [24, 26, 29]. The major changes observed in the transporter expressions during IBD conditions is their down-regulation. These findings suggest that regulating the ABC transporter may allow for further management and treatment for IBD. Similarly, in UC phenotype, the Cysteine and methionine metabolism pathway was enriched. Increased excretion of amino acids such as cysteine and methionine has been used to explain why pediatric IBD patients metabolise protein differently than healthy patients [22].

We observed that multiple genes from species under each cluster are collectively enriched in several pathways in CD and UC across datasets as shown in Figure 3. *ABC transporter* and *Two-component system* pathways were only consistently enriched in CD. ABC transporter, as previously noted, have described their alterations in IBD pathophysiology through multiple studies. Two-component systems (TCSs) pathway is one of the signaling mechanisms, ubiquitously presenting in bacteria and involved in transcriptional regulatory activities such as metabolism and chemotaxis [28, 31]. Shwa et al. described that two-component systems regulate bacterial virulence in response to the interaction between host and the transcriptional regulation of bacterial genes in the gut [28]. The sulfur relay system pathway was enriched in both CD and UC. The association between the sulfur relay system (map04122) pathway and IBD groups (both CD and UC) was found in several studies [16, 23].

Ni et al. observed that the sulfur relay system provides sulfur for biosynthesis of Molybdenum cofactor (Moco) and thiamin, which are sulfur-containing cofactors [23]. As seen in Figure 4, various genes are involved in the synthesis of molybdenum cofactor, catalyzing redox reactions in the bacterial metabolism of sulfur. In Table 7, we found that no single bacterial species has all 29 KOs as annotated by KEGG for sulfur relay systems pathway. This highlights the need for microbial community to work together to perform a common function. Our findings suggest that combining co-occurring species network with pathway enrichment analysis can identify signature pathways and underlying microbial community associated with certain health conditions.

**Table 6:** Result of common enriched metabolic pathways in CD and UC based on Top 5 Hub nodes.

| Phenotype | Enriched pathways based on Top 5 Hub |
| --- | --- |
| CD | Biosynthesis of cofactors (map01240)<br>Cell cycle – Caulobacter (map04112)<br>Homologous recombination (map03440)<br>Mismatch repair (map03430)<br>Protein export (map03060)<br>Pyrimidine metabolism (map00240)<br>Streptomycin biosynthesis (map00521) |
| UC | 2-Oxocarboxylic acid metabolism (map01210)<br>Cell cycle – Caulobacter (map04112)<br>Drug metabolism - other enzymes (map00983)<br>Homologous recombination (map03440)<br>Lysine biosynthesis (map00300)<br>Mismatch repair (map03430)<br>One carbon pool by folate (map00670)<br>Peptidoglycan biosynthesis (map00550) |

**Table 7:** Result of common enriched metabolic pathways in CD and UC based on Top 5 Betweenness nodes.

| Phenotype | Enriched pathways based on Top 5 Betweenness |
| --- | --- |
| CD | None |
| UC | Homologous recombination (map03440)<br>Mismatch repair (map03430)<br>Protein export (map03060)<br>Valine, leucine and isoleucine biosynthesis (map00290) |

Measuring network centrality allows researchers to identify key species that may have important roles within the microbial community. As seen in Table 4, we were able to identify several consistently enriched pathways in CD and UC for both datasets. Even though the top 5 list of CD hub-species for dataset 1 is different from dataset 2, these datasets share commonly enriched pathways. We see parallel patterns with the top 5 list of UC hub-species in datasets 1 and 2, where the datasets share commonly enriched pathways. Interestingly, we found pyrimidine metabolism and streptomycin biosynthesis only appears in CD. Fernandes et al. found that pyrimidine metabolism was significantly enriched in CD and performed an additional validation to identify individual metabolic pathways that are associated with CD through untargeted metabolomics approach [9]. We also measured top 5 list of species with betweenness centrality. We found that valine, leucine and isoleucine biosynthesis is particularly enriched in UC species that have high betweenness centrality. Jagt et al. found that the most differentiating features were increased levels of valine and leucine in IBD versus non-IBD controls [14]. Specifically, the authors observed that the abundance of valine was significantly higher in UC patients.

Our work provides a way to bolster the extraction of microbial species co-occurring analysis with pathway analysis. We augmented microbiome datasets from independent IBD studies to identify co-occurring species in a reliable way. Although the pipeline is agnostic to the number of samples, we utilized case study included a bigger dataset as well as a medium size one. The number of samples between the case versus and control is somehow unbalanced, however the proposed pipeline is able to identify key functional pathways for case vs control comparison within a given dataset and also across datasets, which in some way does overcome the unbalanced number of samples across and within datset.

Therefore, adding more independent IBD studies would be a natural step for future studies to add robustness to the results we presented. There are a handful of current challenges analyzing the complex systems using metagenome data by itself. The whole metagenome sequencing approach allows researchers to investigate microbial profiles in a sample as a community as well as their potential functional profiles. However, this poses a limitation for the identification of genes expressed in different health conditions. The study can be expanded to include analysis of co-occurring species in metatranscriptome datasets as described in this study. This study is conducted to demonstrate the utility of this computational pipeline with IBD studies yet we plan to generalize this pipeline beyond IBD studies.

Overall, our pipeline captures the phenotype-relevant pathways through multi-granularity network analysis from the metagenome abundance datasets. We highlight the functions of microbial communities that form a complex microbial ecosystem, which is based on the species-level microbial associations. We also introduce a computational approach for microbiome researchers to easy-to-implement by incorporating key steps for performing the functional-level analysis of microbial co-occurring communities.

## 6 CONCLUSION

As we get access to more microbiome-relevant data, we gain more opportunities to establish associations between the profiles of a gut microbiome and its health features. While establishing such associations based only on the composition of the microbiome may not always be possible due to the diverse nature of the species forming microbiomes and the roles they can play, we can establish strong associations via signature pathways.

This study demonstrates the viability of identifying characterizing pathways enriched by high co-occurring microbial species for subjects with specific phenotype. To illustrate how such characterizing pathways can be used as biological signals, we use examples of subjects with CD and UC. We developed a computational pipeline specifically designed and implemented to identify these co-occurring species and their associated pathways in a disease phenotype. Using the developed pipeline, we were able to identify enriched pathways associated with the diseased groups that can be used for classification or early recognition of the diseased groups.

This was made possible by finding highly co-occurring microbial communities that significantly contribute to these signature pathways. We believe that the obtained results are important since they point to a deeper understanding of the roles in the microbial community. The study also suggests that further studies of microbiomes may pave the way to non-invasive ways to diagnose certain health conditions at their early stages as well as better assess the progress of various treatment options.

## 7 CODE AVAILABILITY

Code used for this work is publicly available on GitHub: https://github.com/skimicrobe/GutNetMining.

## ACKNOWLEDGMENTS

This work was completed utilizing the Holland Computing Center of the University of Nebraska.

## Notes

### Competing Interest Statement

The authors have declared no competing interest.

## REFERENCES

[1] Francesco Beghini, Lauren J McIver, Aitor Blanco-Míguez, Leonard Dubois, Francesco Asnicar, Sagun Maharjan, Ana Mailyan, Paolo Manghi, Matthias Scholz, Andrew Maltez Thomas, et al. 2021. Integrating taxonomic, functional, and strain-level profiling of diverse microbial communities with bioBakery 3. elife 10 (2021), e65088.

[2] Anne Chao and Tsung-Jen Shen. 2003. Nonparametric estimation of Shannon’s index of diversity when there are unseen species in sample. Environmental and ecological statistics 10 (2003), 429–443.

[3] Meijun Chen and Guangchuang Yu. 2023. MicrobiomeProfiler: An R/shiny package for microbiome functional enrichment analysis. https://github.com/YuLab-SMU/MicrobiomeProfiler/

[4] Paul I Costea, Falk Hildebrand, Manimozhiyan Arumugam, Fredrik Bäckhed, Martin J Blaser, Frederic D Bushman, Willem M De Vos, S Dusko Ehrlich, Claire M Fraser, Masahira Hattori, et al. 2018. Enterotypes in the landscape of gut microbial community composition. Nature microbiology 3, 1 (2018), 8–16.

[5] Gabor Csardi, Tamas Nepusz, et al. 2006. The igraph software package for complex network research. InterJournal, complex systems 1695, 5 (2006), 1–9.

[6] Maria Gloria Dominguez-Bello, Filipa Godoy-Vitorino, Rob Knight, and Martin J Blaser. 2019. Role of the microbiome in human development. Gut 68, 6 (2019), 1108–1114.

[7] Morten Ejby, Folmer Fredslund, Joakim Mark Andersen, Andreja Vujičić Žagar, Jonas Rosager Henriksen, Thomas Lars Andersen, Birte Svensson, Dirk Jan Slotboom, and Maher Abou Hachem. 2016. An ATP binding cassette transporter mediates the uptake of α-(1, 6)-linked dietary oligosaccharides in Bifidobacterium and correlates with competitive growth on these substrates. Journal of Biological Chemistry 291, 38 (2016), 20220–20231.

[8] Kai Feng, Xi Peng, Zheng Zhang, Songsong Gu, Qing He, Wenli Shen, Zhujun Wang, Danrui Wang, Qiulong Hu, Yan Li, et al. 2022. iNAP: an integrated network analysis pipeline for microbiome studies. iMeta 1, 2 (2022), e13.

[9] Philip Fernandes, Yash Sharma, Fatima Zulqarnain, Brooklyn McGrew, Aman Shrivastava, Lubaina Ehsan, Dawson Payne, Lillian Dillard, Deborah Powers, Isabelle Aldridge, et al. 2023. Identifying metabolic shifts in Crohn’s disease using’omics-driven contextualized computational metabolic network models. Scientific Reports 13, 1 (2023), 203.

[10] Eric A Franzosa, Alexandra Sirota-Madi, Julian Avila-Pacheco, Nadine Fornelos, Henry J Haiser, Stefan Reinker, Tommi Vatanen, A Brantley Hall, Himel Mallick, Lauren J McIver, et al. 2019. Gut microbiome structure and metabolic activity in inflammatory bowel disease. Nature microbiology 4, 2 (2019), 293–305.

[11] Daniel M Griffith, Joseph A Veech, and Charles J Marsh. 2016. Cooccur: probabilistic species co-occurrence analysis in R. Journal of Statistical Software 69 (2016), 1–17.

[12] Kaijian Hou, Zhuo-Xun Wu, Xuan-Yu Chen, Jing-Quan Wang, Dongya Zhang, Chuanxing Xiao, Dan Zhu, Jagadish B Koya, Liuya Wei, Jilin Li, et al. 2022. Microbiota in health and diseases. Signal transduction and targeted therapy 7, 1 (2022), 135.

[13] Ying Huang, Pascale Anderle, Kimberly J Bussey, Catalin Barbacioru, Uma Shankavaram, Zunyan Dai, William C Reinhold, Audrey Papp, John N Weinstein, and Wolfgang Sadée. 2004. Membrane transporters and channels: role of the transportome in cancer chemosensitivity and chemoresistance. Cancer research 64, 12 (2004), 4294–4301.

[14] Jasmijn Z Jagt, Eduard A Struys, Ibrahim Ayada, Abdellatif Bakkali, Erwin EW Jansen, Jürgen Claesen, Johan E van Limbergen, Marc A Benninga, Nanne KH de Boer, and Tim GJ de Meij. 2022. Fecal amino acid analysis in newly diagnosed pediatric inflammatory bowel disease: a multicenter case-control study. Inflammatory Bowel Diseases 28, 5 (2022), 755–763.

[15] Zachary D Kurtz, Christian L Müller, Emily R Miraldi, Dan R Littman, Martin J Blaser, and Richard A Bonneau. 2015. Sparse and compositionally robust inference of microbial ecological networks. PLoS computational biology 11, 5 (2015), e1004226.

[16] James D Lewis, Eric Z Chen, Robert N Baldassano, Anthony R Otley, Anne M Griffiths, Dale Lee, Kyle Bittinger, Aubrey Bailey, Elliot S Friedman, Christian Hoffmann, et al. 2015. Inflammation, antibiotics, and diet as environmental stressors of the gut microbiome in pediatric Crohn’s disease. Cell host & microbe 18, 4 (2015), 489–500.

[17] Dong Li, Haowei Ni, Shuo Jiao, Yahai Lu, Jizhong Zhou, Bo Sun, and Yuting Liang. 2021. Coexistence patterns of soil methanogens are closely tied to methane generation and community assembly in rice paddies. Microbiome 9 (2021), 1–13.

[18] Jason Lloyd-Price, Cesar Arze, Ashwin N Ananthakrishnan, Melanie Schirmer, Julian Avila-Pacheco, Tiffany W Poon, Elizabeth Andrews, Nadim J Ajami, Kevin S Bonham, Colin J Brislawn, et al. 2019. Multi-omics of the gut microbial ecosystem in inflammatory bowel diseases. Nature 569, 7758 (2019), 655–662.

[19] Weijun Luo. 2017. Pathview: pathway based data integration and visualization. Bioconductor Documentation p-8 (2017).

[20] Bin Ma, Yiling Wang, Shudi Ye, Shan Liu, Erinne Stirling, Jack A Gilbert, Karoline Faust, Rob Knight, Janet K Jansson, Cesar Cardona, et al. 2020. Earth microbial co-occurrence network reveals interconnection pattern across microbiomes. Microbiome 8 (2020), 1–12.

[21] Fabien Magne, Martin Gotteland, Lea Gauthier, Alejandra Zazueta, Susana Pesoa, Paola Navarrete, and Ramadass Balamurugan. 2020. The firmicutes/bacteroidetes ratio: a relevant marker of gut dysbiosis in obese patients? Nutrients 12, 5 (2020), 1474.

[22] Francois-Pierre Martin, Ming-Ming Su, Guo-Xiang Xie, Seu Ping Guiraud, Martin Kussmann, Jean-Philippe Godin, Wei Jia, and Andreas Nydegger. 2017. Urinary metabolic insights into host-gut microbial interactions in healthy and IBD children. World journal of gastroenterology 23, 20 (2017), 3643.

[23] Josephine Ni, Ting-Chin David Shen, Eric Z Chen, Kyle Bittinger, Aubrey Bailey, Manuela Roggiani, Alexandra Sirota-Madi, Elliot S Friedman, Lillian Chau, Andrew Lin, et al. 2017. A role for bacterial urease in gut dysbiosis and Crohn’s disease. Science translational medicine 9, 416 (2017), eaah6888.

[24] V Nirmal and K Reena. 2018. ATP-binding cassette (ABC) transporters and their role in inflammatory bowel disease (IBD). Biomed J Sci Tech Res 5 (2018).

[25] Jari Oksanen, FG Blanchet, R Kindt, PMPR Legendre, PR Minchin, R. O’hara, G Simpson, P Solymos, M Henry, H Stevens, et al. 2017. Ordination methods, diversity analysis and other functions for community and vegetation ecologists. Vegan: Community Ecol Package (2017), 05–26.

[26] Sandra Pérez-Torras, Ingrid Iglesias, Marta Llopis, Juan J Lozano, María Antolín, Francisco Guarner, and Marçal Pastor-Anglada. 2016. Transportome profiling identifies profound alterations in Crohn’s disease partially restored by commensal bacteria. Journal of Crohn’s and Colitis 10, 7 (2016), 850–859.

[27] Stefanie Peschel, Christian L Müller, Erika Von Mutius, Anne-Laure Boulesteix, and Martin Depner. 2021. NetCoMi: network construction and comparison for microbiome data in R. Briefings in bioinformatics 22, 4 (2021), bbaa290.

[28] Claire Shaw, Matthias Hess, and Bart C Weimer. 2022. Two-component systems regulate bacterial virulence in response to the host gastrointestinal environment and metabolic cues. Virulence 13, 1 (2022), 1666–1680.

[29] Nirmal Verma, Vineet Ahuja, and Jaishree Paul. 2011. Mechanism of dysregulation of ABCF2 transporter in Ulcerative colitis patients: P-244. Inflammatory Bowel Diseases 17 (2011), S84.

[30] Aleksej Zelezniak, Sergej Andrejev, Olga Ponomarova, Daniel R Mende, Peer Bork, and Kiran Raosaheb Patil. 2015. Metabolic dependencies drive species co-occurrence in diverse microbial communities. Proceedings of the National Academy of Sciences 112, 20 (2015), 6449–6454.

[31] Yitian Zhou, Qinqin Pu, Jiandong Chen, Guijuan Hao, Rong Gao, Afsar Ali, Ansel Hsiao, Ann M Stock, Mark Goulian, and Jun Zhu. 2021. Thiol-based functional mimicry of phosphorylation of the two-component system response regulator ArcA promotes pathogenesis in enteric pathogens. Cell reports 37, 12 (2021), 110147.

